## Supplementary material for "Birds migrate longitudinally in response to the resultant Asian monsoons of the Qinghai-Tibet Plateau uplift": Figures S1-S11 Table S1

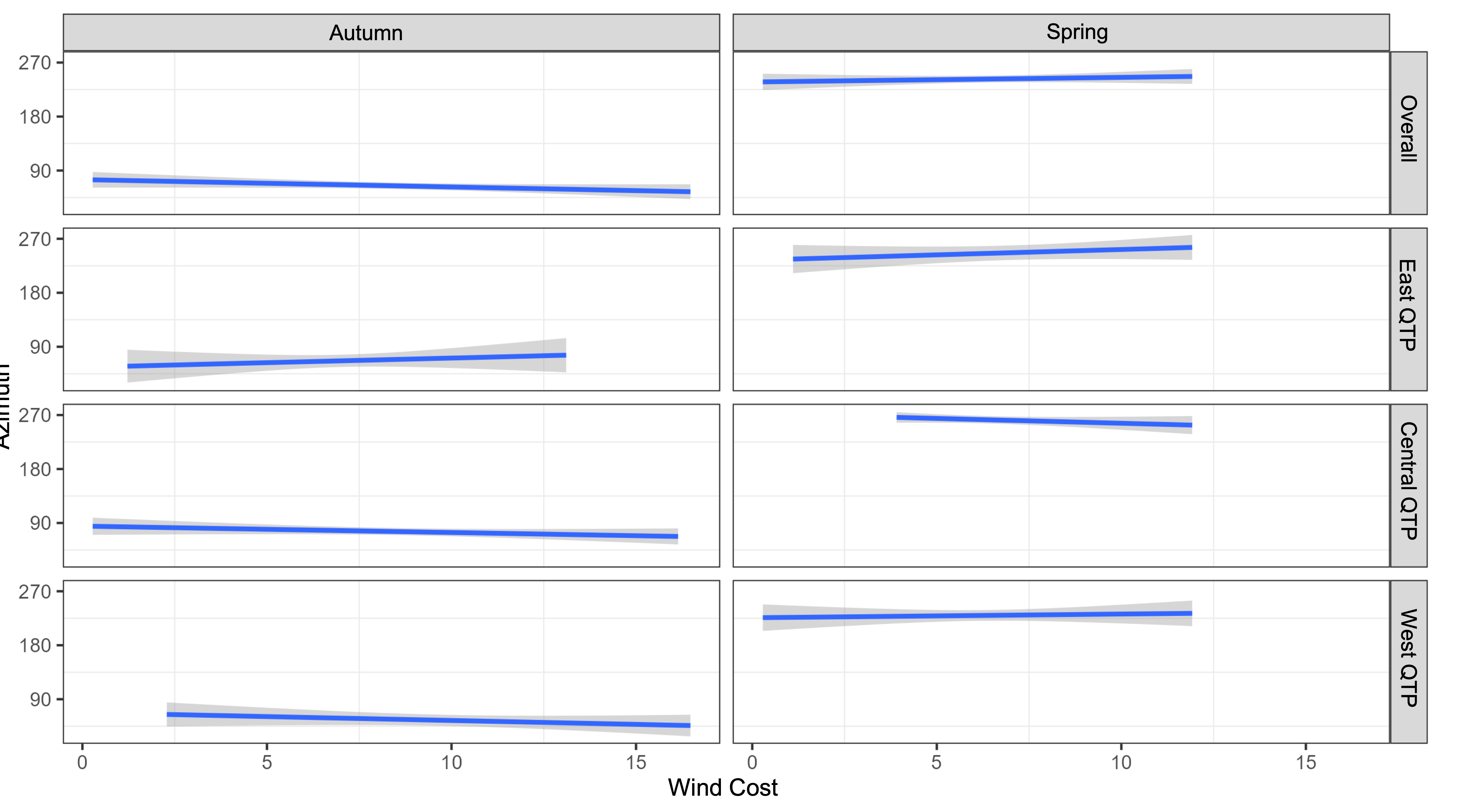

**Fig. S1.** **Migration azimuth changes with wind cost in the area west, central, and east of the Qinghai-Tibet Plateau during studied avian migration periods**. Wind represents the wind cost calculated by wind connectivity. West QP, Central QTP, and East QTP denote areas west (longitude < 73°E), central (73°E ≤ longitude < 105°E), and east (longitude ≥ 105°E) of the Qinghai-Tibet Plateau, respectively.

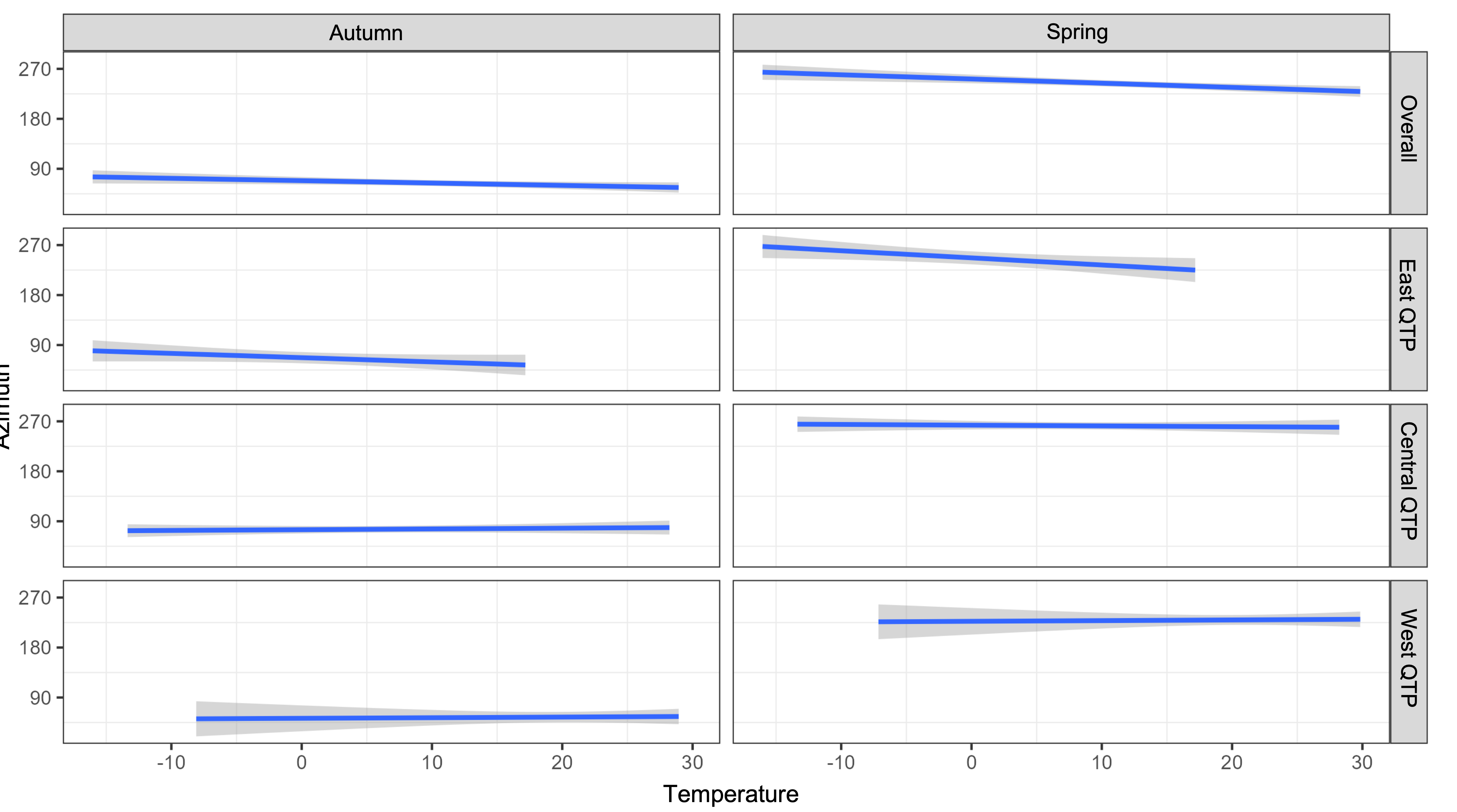
Fig. S2. Migration azimuth changes with average annual temperature in the area west, central, and east of the Qinghai-Tibet Plateau during studied avian migration periods. West QTP, Central QTP, and East QTP denote areas west (longitude < 73°E), central (73°E ≤ longitude < 105°E), and east (longitude ≥ 105°E) of the Qinghai-Tibet Plateau, respectively.

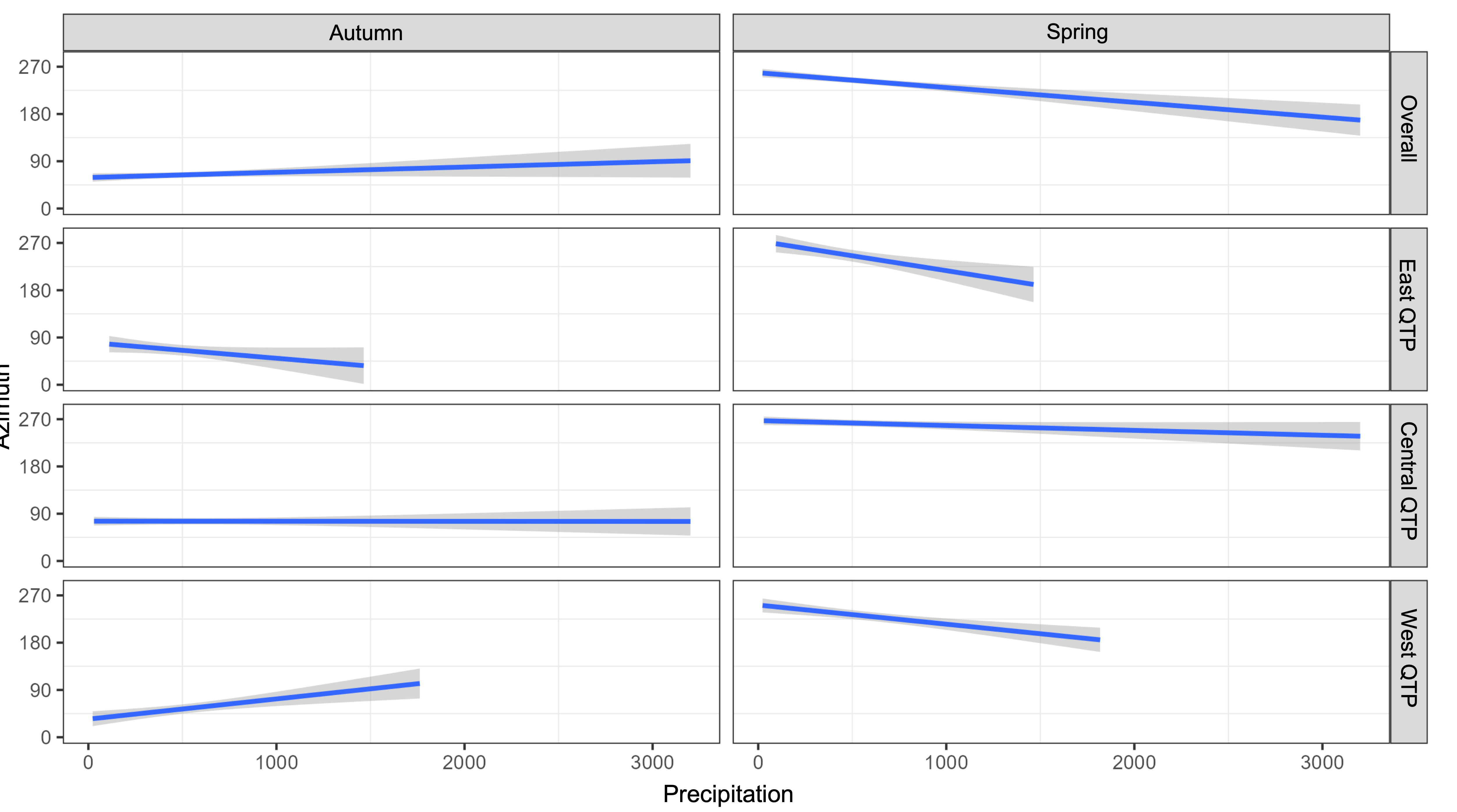

**Fig. S3. Migration azimuth changes with average annual precipitation in the area west, central, and east of the Qinghai-Tibet Plateau during studied avian migration periods.** West QTP, Central QTP, and East QTP denote areas west (longitude < 73°E), central (73°E ≤ longitude < 105°E), and east (longitude ≥ 105°E) of the Qinghai-Tibet Plateau, respectively.

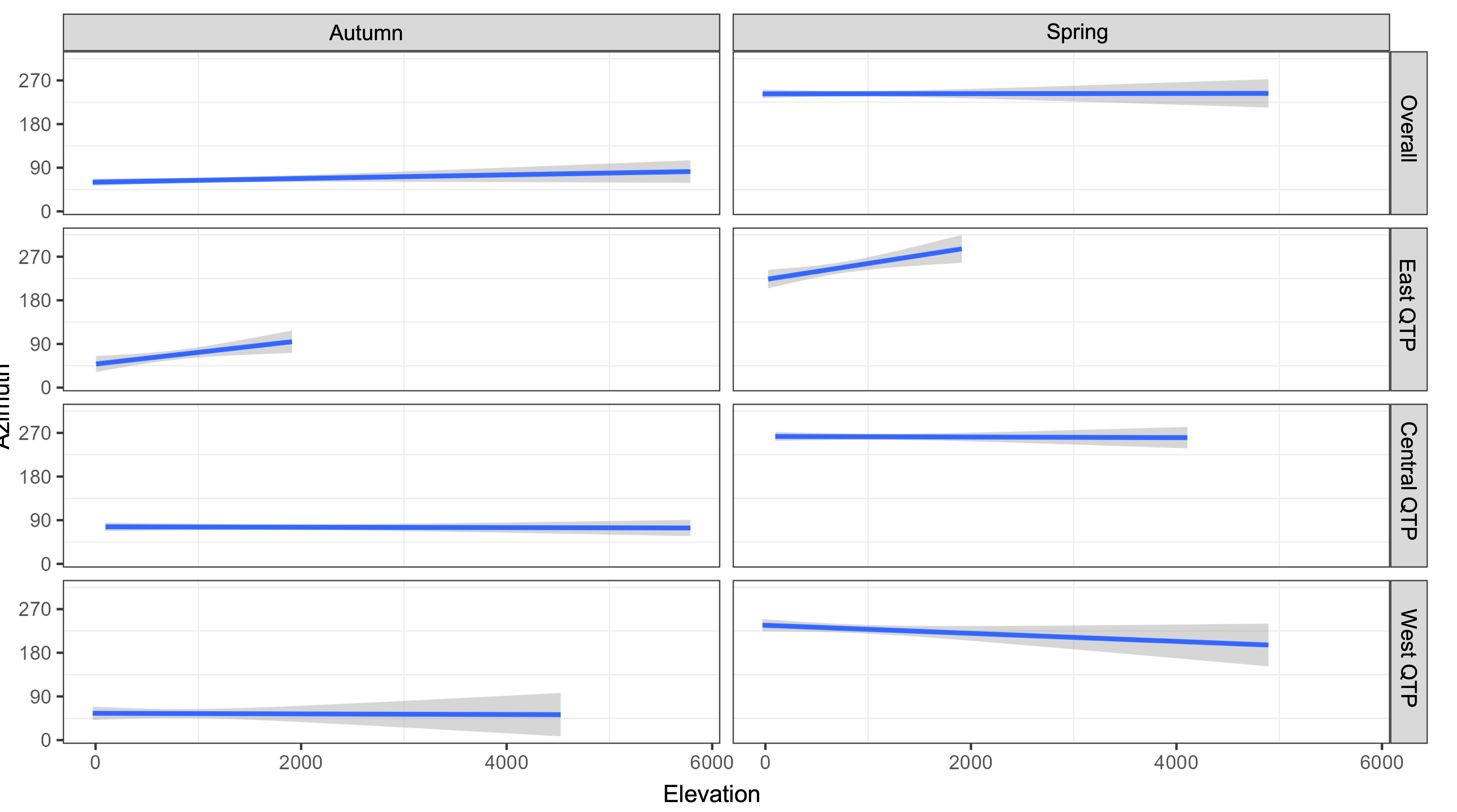
**Fig. S4. Migration azimuth changes with altitude in the west, central, and east of the Qinghai-Tibet Plateau during studied avian migration periods**. West QTP, central QTP, and East QTP denote areas west (longitude < 73°E), central (73°E ≤ longitude < 105°E), and east (longitude ≥ 105°E) of the Qinghai-Tibet Plateau, respectively.

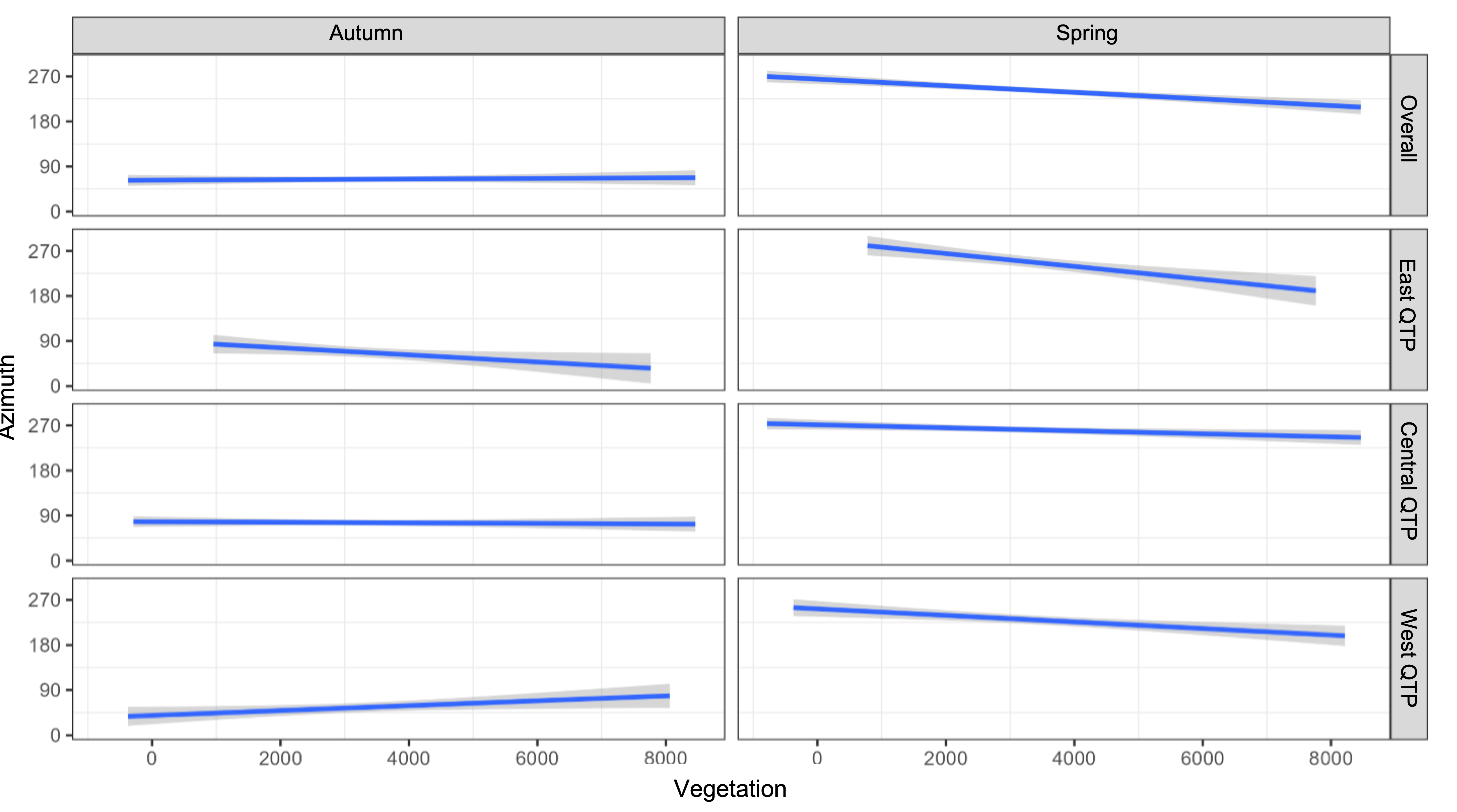
**Fig. S5. Migration azimuth changes with measured vegetation in the west, central, and east of the Qinghai-Tibet Plateau during studied avian migration periods.** Vegetation is measured by the proportion of average vegetation cover in each pixel. West QTP, Central QTP, and East QTP denote areas west (longitude < 73°E), central (73°E ≤ longitude < 105°E), and east (longitude ≥ 105°E) of the Qinghai-Tibet Plateau, respectively.

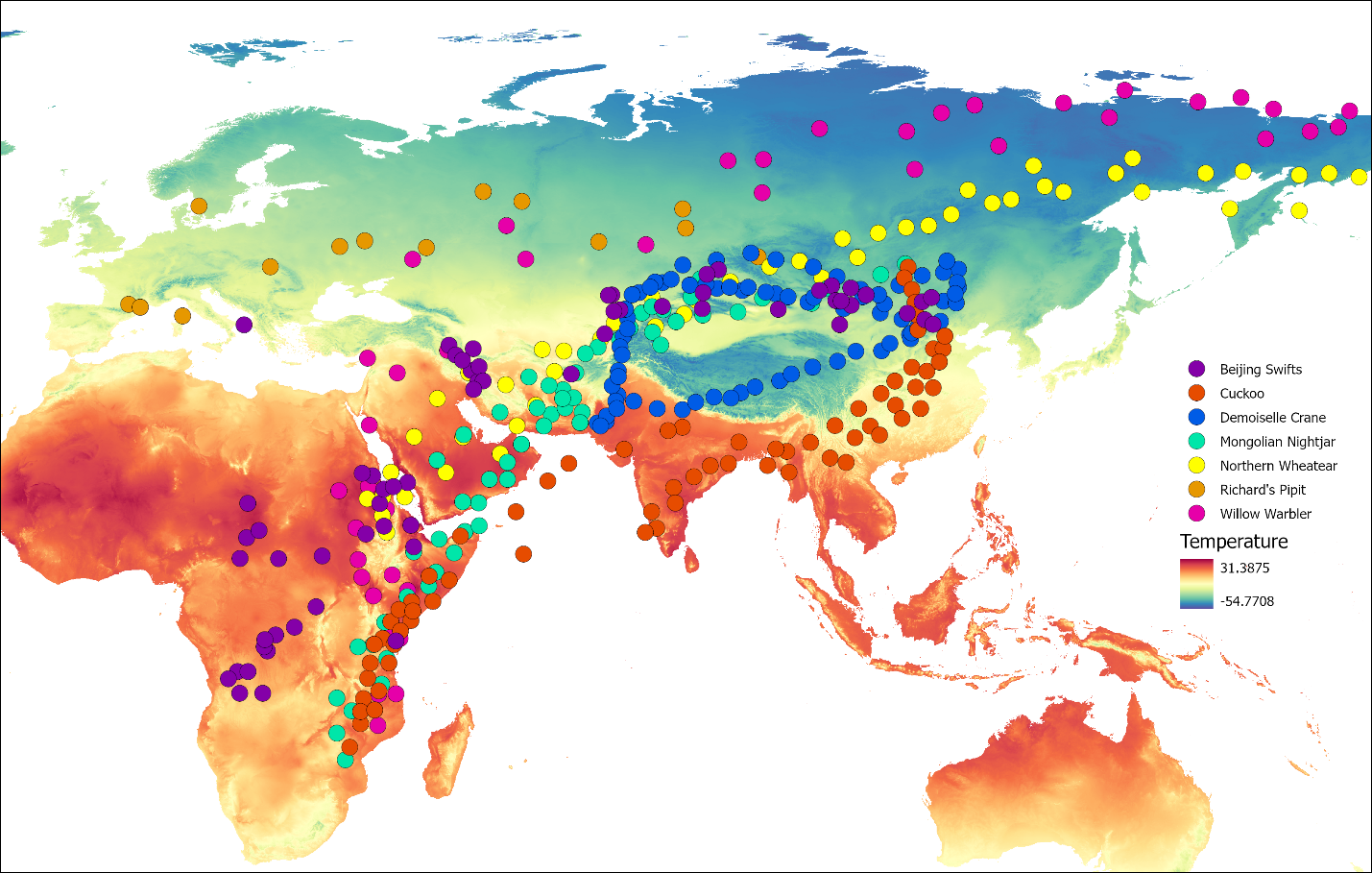

**Fig. S6. Birds migrate along with the gradient of annual temperature in the study area.** Round points with different colours represent tracking records of the seven species that migrate across the Qinghai-Tibet Plateau.

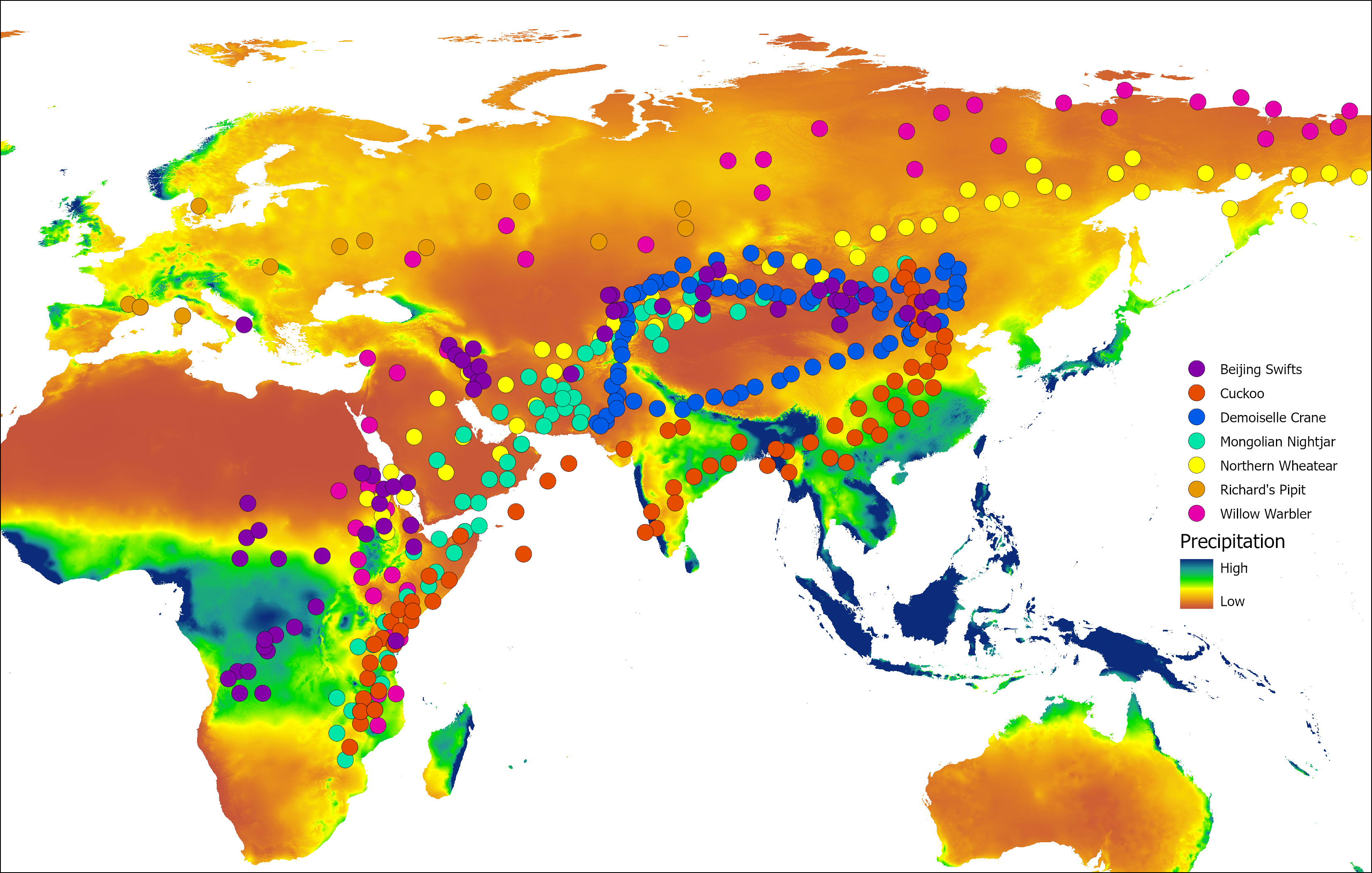

**Fig. S7. Birds migrate along with the precipitation gradient.** Round points with different colours represent tracking records of multiple species that migrate across the Qinghai-Tibet Plateau.

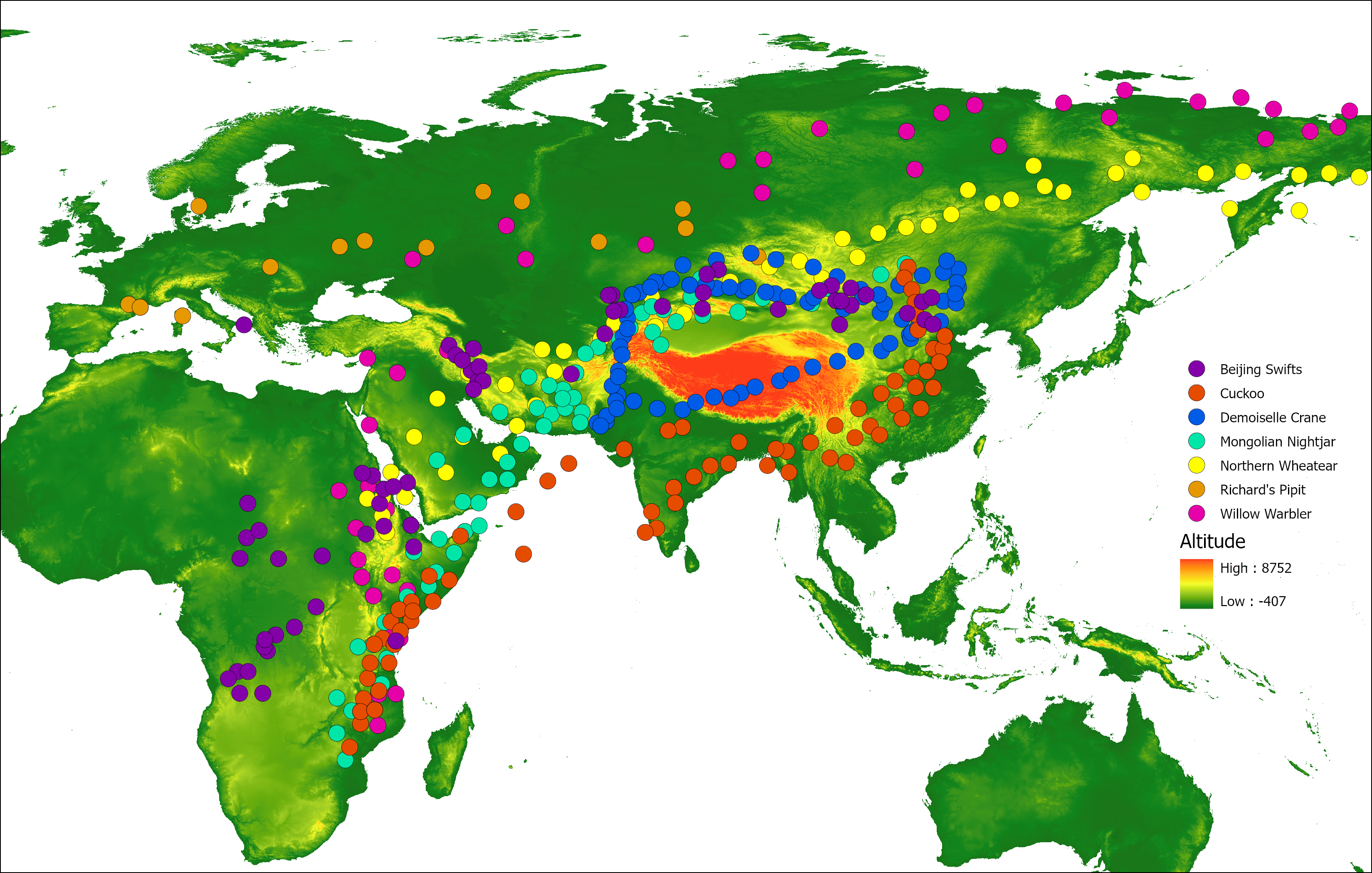

**Fig. S8. Birds migrate along with the vegetation gradient.** Round points with different colours represent tracking records of multiple species that migrate across the Qinghai-Tibet Plateau.

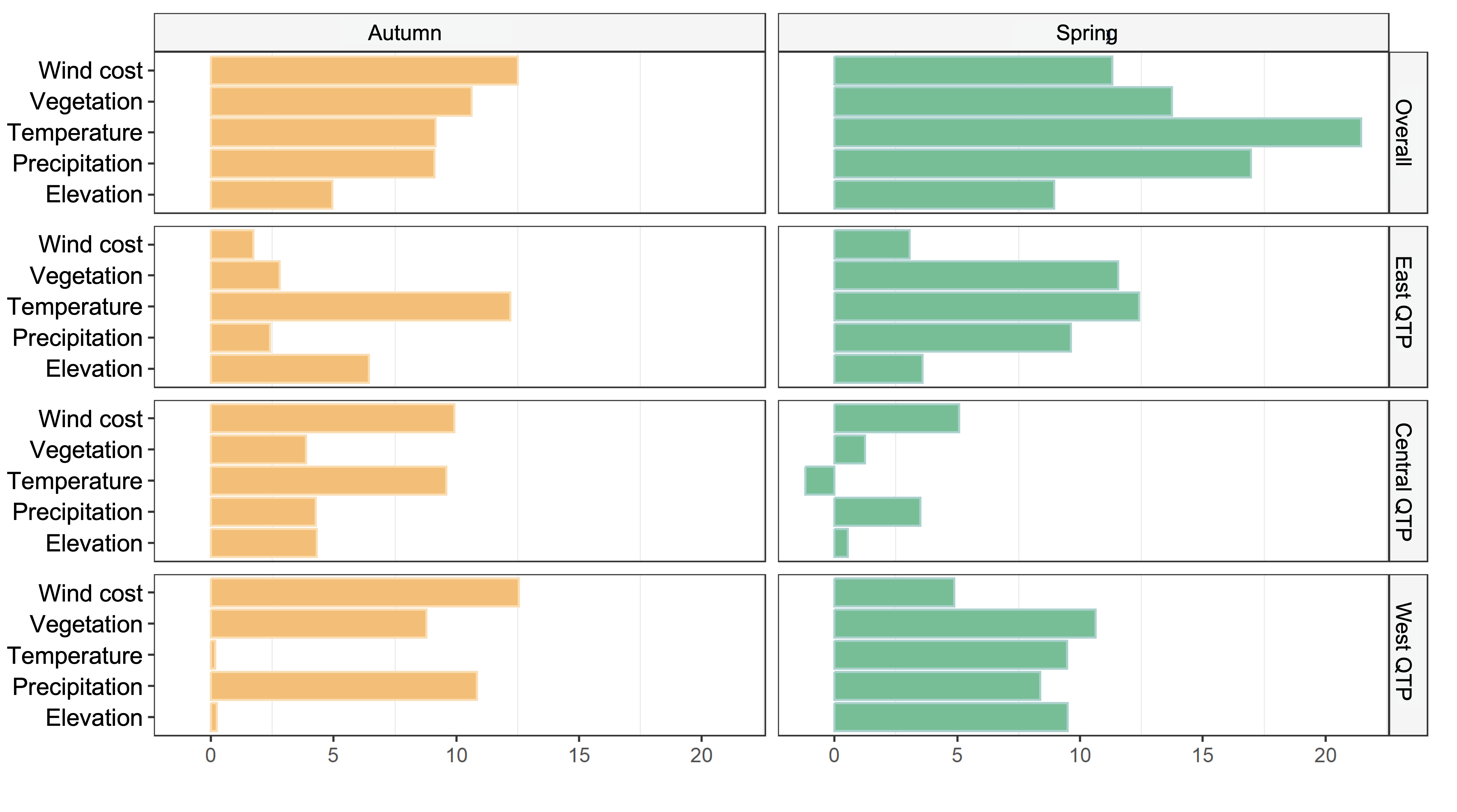

**Fig. S9 Environmental factors that influence avian migratory strategies in the west, central, and east of the Qinghai-Tibet Plateau.** Using Random Forest to model the correlation between environmental factors and migratory strategies (migration direction) along different stages of migration, the bar shows how model performance is reduced by removing the corresponding factor. Wind represents the wind cost calculated by wind connectivity. Vegetation is measured by the proportion of average vegetation cover in each pixel. Temperature is the average annual temperature. Precipitation is the average annual precipitation. West QTP, Central QTP, and East QTP denote areas west (longitude < 73°E), central (73°E ≤ longitude < 105°E), and east of (longitude ≥ 105°E) the Qinghai-Tibet Plateau, respectively.

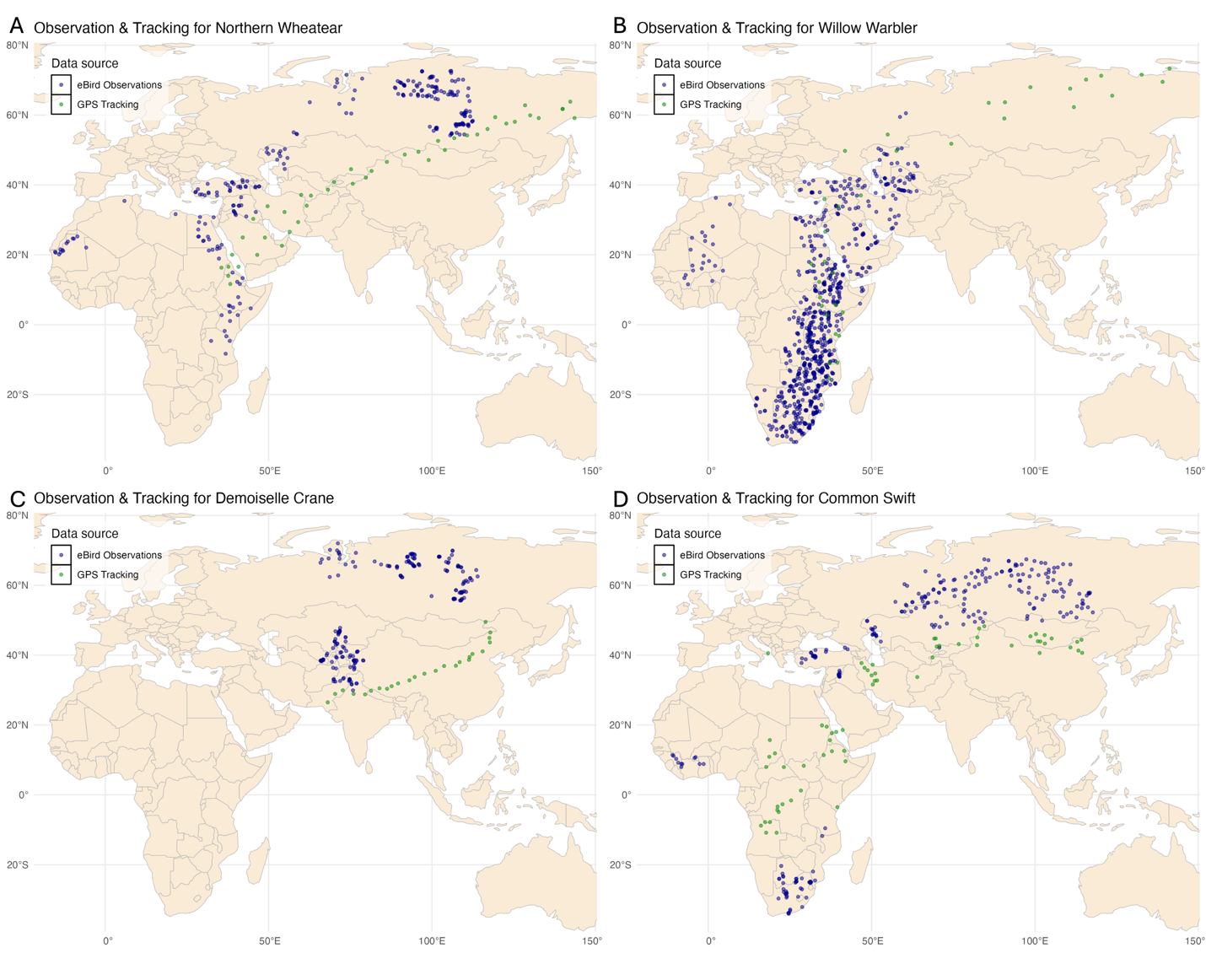

**Fig. S10. Distribution of eBird and GPS tracking data for Northern Wheatear, Willow Warbler, Demoiselle Crane and Common Swift.** Blue points denote eBird observations and green points denote GPS tracking data.

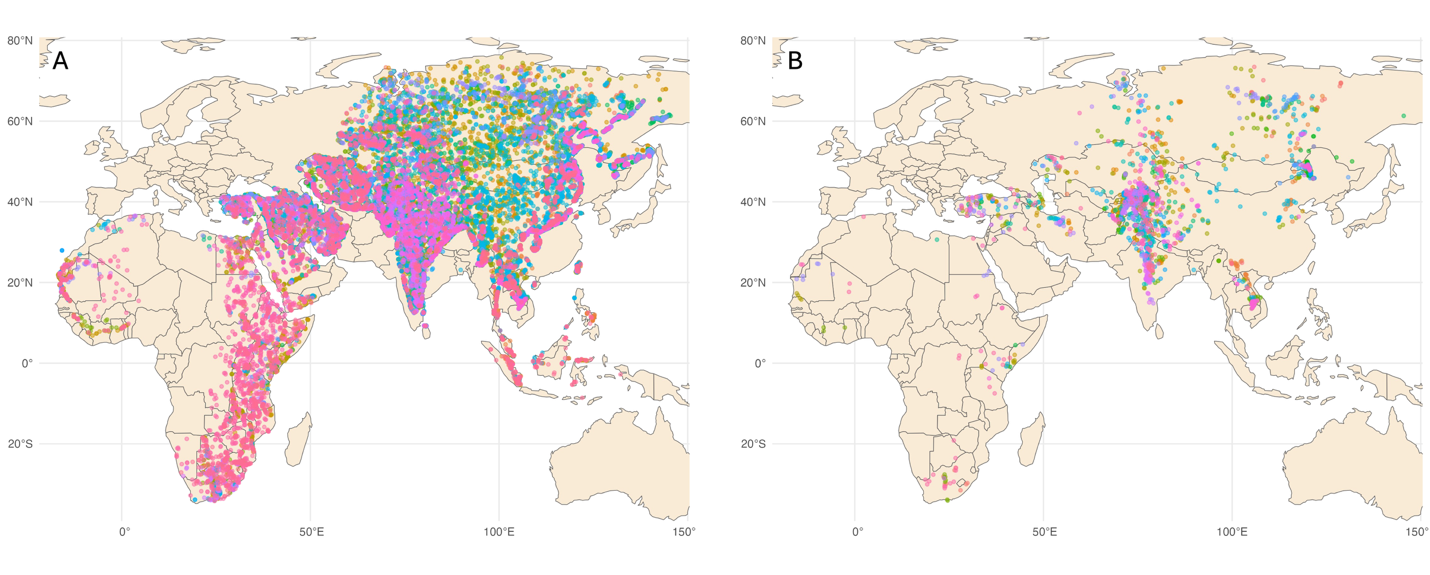

**Figure S11. Distribution of eBird observations for all 50 species modified by our weekly dynamic model.** (A) shows all possible occurrence for each of the 50 species and (B) shows the most possible occurrence of each species in each migration week. We derived the occurrence for each species using the Adaptive Spatio-Temporal Model (AdaSTEM) that we developed (see Methods for details). Different colours represent different species.

Table S1. List of species analysed in our study and AUC values for modelling their distributions.

|  | **Species** | **Latin** | **AUC (Breeding)** | **AUC (Wintering)** |
| --- | --- | --- | --- | --- |
| **1** | **Amur Falcon*** | *Falco amurensis* | 0.9887 | 0.9915 |
| **2** | Bar-headed Goose | *Anser indicus* | 0.9097 | 0.9277 |
| **3** | Barn Swallow | *Hirundo rustica* | 0.962 | 0.97 |
| **4** | **Black Kite*** | *Milvus migrans* | - | - |
| **5** | **Black-headed Gull*** | *Chroicocephalus ridibundus* | - | - |
| **6** | **Black-tailed godwit*** | *Limosa limosa* | - | - |
| **7** | Bluethroat | *Cyanecula svecica* | 0.9741 | 0.9627 |
| **8** | Booted Eagle | *Hieraaetus pennatus* | 0.9693 | 0.9757 |
| **9** | Brown Shrike | *Lanius cristatus* | 0.9704 | 0.9634 |
| **10** | Citrine Wagtail | *Motacilla citreola* | 0.9538 | 0.9669 |
| **11** | **Common Crane*** | *Grus grus* | 0.9951 | 0.9755 |
| **12** | **Common Cuckoo*** | *Cuculus canorus* | 0.9831 | 0.9643 |
| **13** | Common Greenshank | *Tringa nebularia* | 0.9212 | 0.9228 |
| **14** | **Common Redshank*** | *Tringa totanus* | 0.9241 | 0.9476 |
| **15** | Common Ringed Plover | *Charadrius hiaticula* | 0.9845 | 0.967 |
| **16** | Common Rosefinch | *Carpodacus erythrinus* | 0.9543 | 0.9775 |
| **17** | Common Snipe | *Gallinago gallinago* | 0.992 | 0.9404 |
| **18** | **Common Swift*** | *Apus apus* | 0.9578 | 0.9723 |
| **19** | Curlew Sandpiper | *Calidris ferruginea* | 0.9633 | 0.9782 |
| **20** | **Demoiselle Crane*** | *Anthropoides virgo* | 0.9846 | 0.9871 |
| **21** | Dunlin | *Calidris alpina* | 0.9852 | 0.9394 |
| **22** | Eurasian Hobby | *Falco subbuteo* | 0.9584 | 0.9934 |
| **23** | **Eurasian Nightjar*** | *Caprimulgus europaeus* | 0.9802 | 0.9888 |
| **24** | Eurasian Whimbrel | *Numenius phaeopus* | 0.9528 | 0.9347 |
| **25** | Eurasian Wryneck | *Jynx torquilla* | 0.9625 | 0.9529 |
| **26** | Eurasian Wigeon | *Mareca penelope* | 0.9704 | 0.9551 |
| **27** | Gadwall | *Mareca strepera* | 0.9691 | 0.9482 |
| **28** | Greater Short-toed Lark | *Calandrella brachydactyla* | 0.9574 | 0.983 |
| **29** | Greater Spotted Eagle | *Clanga clanga* | 0.9431 | 0.9281 |
| **30** | Greenish Warbler | *Phylloscopus trochiloides* | 0.9548 | 0.9769 |
| **31** | Kentish Plover | *Charadrius alexandrinus* | 0.9388 | 0.9417 |
| **32** | Lesser Kestrel | *Falco naumanni* | 0.9642 | 0.9102 |
| **33** | Little Bunting | *Emberiza pusilla* | 0.9951 | 0.9614 |
| **34** | Little Ringed Plover | *Charadrius dubius* | 0.9248 | 0.9477 |
| **35** | Little Stint | *Calidris minuta* | 0.9647 | 0.9611 |
| **36** | Long-legged Buzzard | *Buteo rufinus* | 0.973 | 0.9744 |
| **37** | Marsh Sandpiper | *Tringa stagnatilis* | 0.9404 | 0.9388 |
| **38** | **Montagu's harrier*** | *Circus pygargus* | - | - |
| **39** | Northern Lapwing | *Vanellus vanellus* | 0.9791 | 0.9358 |
| **40** | Northern Shoveler | *Spatula clypeata* | 0.9544 | 0.9419 |
| **41** | **Northern Wheatear*** | *Oenanthe oenanthe* | 0.979 | 0.9853 |
| **42** | **Osprey*** | *Pandion haliaetus* | - | - |
| **43** | **Pallid Harrier*** | *Circus macrourus* | 0.9928 | 0.9838 |
| **44** | Pied Avocet | *Recurvirostra avosetta* | 0.9571 | 0.961 |
| **45** | **Peregrine Falcons*** | *Falco peregrinus* | - | - |
| **46** | **Richard's pipit*** | *Anthus richardi* | 0.9113 | 0.9356 |
| **47** | Ruff | *Calidris pugnax* | 0.982 | 0.958 |
| **48** | Short-toed Snake-Eagle | *Circaetus gallicus* | 0.9707 | 0.9857 |
| **49** | **Siberian Gull*** | *Larus fuscus heuglini* | - | - |
| **50** | **Siberian Rubythroat*** | *Calliope calliope* | - | - |
| **51** | Spotted Redshank | *Tringa erythropus* | 0.9675 | 0.9688 |
| **52** | Steppe Eagle | *Aquila nipalensis* | 0.9973 | 0.9648 |
| **53** | Terek Sandpiper | *Xenus cinereus* | 0.9626 | 0.9737 |
| **54** | Tree Pipit | *Anthus trivialis* | 0.9774 | 0.9752 |
| **55** | Western Yellow Wagtail | *Motacilla flava* | 0.9609 | 0.9536 |
| **56** | **Willow Warbler*** | *Phylloscopus trochilus* | 0.9768 | 0.9808 |
| **57** | Wood Sandpiper | *Tringa glareola* | 0.9256 | 0.9283 |
| **58** | Yellow-breasted Bunting | *Emberiza aureola* | 0.9959 | 0.9969 |

* Tracking data are available. – eBird observations are not large enough to support our model.
